# Envelope analysis of the human alpha rhythm reveals EEG Gaussianity

**DOI:** 10.1101/2021.03.31.437785

**Authors:** Víctor Manuel Hidalgo, Javier Díaz, Jorge Mpodozis, Juan-Carlos Letelier

## Abstract

The origin of the human alpha rhythm has been a matter of debate since Lord Adrian attributed it to synchronous neural populations in the occipital cortex. Although some authors have pointed out the Gaussian characteristics of the alpha rhythm, their results have been repeatedly disregarded in favor of Adrian’s interpretation; even though the first EEG Gaussianity reports can be traced back to the origins of the field. Here we revisit this problem using the envelope analysis — a method that relies on the fact that the coefficient of variation of the envelope (CVE) for continuous-time zero-mean Gaussian noise (as well as for any filtered sub-band) is equal to 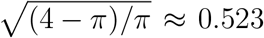, thus making the CVE a fingerprint for Gaussianity. As a consequence, any significant deviation from 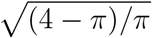 is linked to synchronous neural dynamics. We analyzed occipital EEG and iEEG data from massive public databases. Our results showed the human alpha rhythm can be characterized either as a synchronous or as a Gaussian signal based on the value of its CVE. Furthermore, Fourier analysis showed the canonical spectral peak at ≈ 10[Hz] is present in both the synchronous and Gaussian cases, thus demonstrating this same peak can be produced by different underlying neural dynamics. This study confirms the original interpretation of Adrian regarding the origin of the alpha rhythm but also opens the door for the study of Gaussianity in brain dynamics. These results suggest a broader interpretation for event-related synchronization/desynchronization (ERS/ERD) may be needed. Envelope analysis constitutes a novel complement to Fourier-based methods for neural signal analysis relating amplitude modulation patterns (CVE) to signal energy.

## 1 Introduction

The neuronal dynamics that generate the human alpha rhythm, first described by Berger (1929), have been a matter for discussion since Lord Adrian attributed it to synchronous neural populations in the occipital cortex. Adrian undertook a thorough examination of the alpha rhythm and attributed its origin to neural synchronization (Adrian and Matthews, 1934); an idea that rapidly became mainstream and accepted as the standard interpretation for EEG phenomena. Indeed, neural synchronization is a fundamental mechanism in EEG signal generation (Buzsáki et al., 2012) and a widespread phenomenon at many levels of biological organization (Singer, 1999; Varela et al., 2001). Adrian also reported that the rhythm can be entrained by a flicker to frequencies as high as 25 [Hz]. The famous cybernetician Norbert Wiener hypothesized that the alpha rhythm is produced by a synchronous population of coupled nonlinear oscillators with similar natural frequencies (Wiener, 1948; Strogatz, 1994) — a claim supported by Kuramoto’s early model of coupled oscillators systems (Kuramoto, 1975).

Nevertheless, some authors have reported that the alpha rhythm exhibits Gaussian characteristics, but this possibility has been consistently disregarded for any EEG phenomena under the assumption that the amplitude of a signal emerging from an asynchronous population is either zero or, at best, too low to be measured (Cohen, 2017; Lachaux et al., 1999; Singer, 1999). On the contrary, the amplitude of a signal emerging from a population of asynchronous oscillators is proportional to 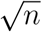 (Elul, 1972), as was recognized early in the EEG literature by Motokawa (1943). In fact, Motokawa contended that the alpha rhythm originated from a population of uncorrelated oscillators (Motokawa and Mita, 1942). In an elegant approach, Saunders (1963) compared the distribution of raw signal amplitude values of “well developed” alpha rhythm epochs against the Gaussian distribution and found a close fit. From this perspective, some researchers have suggested that the underlying neural generator may be better described as an alpha filter (Lopes Da Silva et al., 1973): a system that receives random input and produces the characteristic narrow band signal centered ≈ 10[Hz]. This interpretation of the alpha rhythm generator as a bandpass filter can be traced back to Prast (1949). Dick and Vaughn (1970) went one step further and estimated the envelope function by the complex demodulation method. They found the envelope values of alpha rhythm closely fit the Rayleigh distribution, which is important because the envelope function of a Gaussian signal follows that distribution (Schwartz et al., 1995). Furthermore, they proposed to study the changes of the variance of the envelope to explore the time-variant behaviour of the alpha rhythm with an approach similar to ours, but which did not exploit the properties of the envelope of Gaussian noise.

Another remarkable characteristic of the alpha rhythm is what Adrian and Matthews described as “waxing and waning” or “pulsating” activity, i.e. a distinctive amplitude modulation pattern. While sometimes ignored, there is some interindividual variability among alpha amplitude modulation patterns (Niedermeyer, 2005). Davis and Davis (1936) reported as many as four distinct alpha rhythm types based on their amplitude modulation, including cases where little or no waning is present. These distinct patterns are also reflected in the frequency of the alpha rhythm envelope (Schroeder and Barr, 2000). Consequently, these different alpha modulation regimes result in characteristic raw signal and envelope waveforms. Interestingly, in recent years, some authors have pointed out the importance of the signal waveform to detect features in neural signals that may not be evident using Fourier-based methods (Cole and Voytek, 2017). Indeed, as we will show, the signal waveform does offer information about neural dynamics; information that can be extracted analyzing the envelope signal.

In a previous report, Díaz et al. (2018) used rat EEG to describe a novel framework to characterize EEG signals: the envelope analysis. This analysis relies on the fact that the coefficient of variation of the envelope (CVE) of a continuous-time zero-mean Gaussian noise signal (and any of its sub-filtered bands) is invariant and equal to 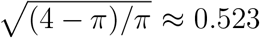 (Schwartz et al., 1995). Thus, the Gaussian CVE is a universal constant and acts as a mark of Gaussianity. To capture this time-variant behaviour, the signal under study was segmented and the CVE for each epoch estimated, as well as the mean of the envelope (ME) to measure energy. As Gaussianity is a hallmark of asynchrony, any deviations from the CVE of Gaussianity reveals synchronous processes. Díaz et al. (2007) showed that a signal emerging from a population of phase-locked nonlinear oscillators exhibits low-CVE values, while high-CVE signals emerged only from unsteady states. Hence, CVE (morphology) and ME (energy) can be combined to produce a novel phase-space in which signals can be categorized: the envelope characterization space. This space unifies generator dynamics, energy and signal morphology in a single framework. Using this framework, we propose here to classify EEG epochs not by frequency alone but by the two parameters defining the envelope characterization space: CVE and the mean energy of its envelope.

## 2 Materials and Methods

### 2.1 Data

#### 2.1.1 EEG data

We applied the envelope analysis to the EEG data of the “Leipzig Study for Mind-Body-Emotion Interactions” (LEMON) dataset (Babayan et al., 2019). This is a cross-sectional, publicly available dataset that contains resting-state EEG data from 216 younger and older healthy adults and is part of the MPI Leipzig MindBrain-Body database. The study protocol was in accordance with the declaration of Helsinki. All participants provided written consent prior to assessments and were screened for drug use, neurological, cardiovascular, and psychiatric conditions.

The recordings were obtained using 62 active electrodes, 61 EEG channels in accordance with the 10-10 system (Oostenveld and Praamstra, 2001) and 1 VEOG electrode below the right eye. The signals were recorded with a bandpass filter between 0.015[Hz] and 1[kHz] and sampled with a frequency of 2500[Hz]. Each recording comprised 16 blocks of 60[s], 8 blocks with eyes closed (EC) and 8 blocks with eyes open (EO). The recording began with eyes closed and blocks were interleaved. The raw data from 13 participants was discarded due to defective metadata and low data quality. Data from the remaining 203 subjects was preprocessed as follows. The data was downsampled from 2500[Hz] to 250[Hz] and bandpass filtered between 1-45[Hz] with an 8th order Butterworth filter. Blocks were split into EC and EO conditions for further processing. Outlier channels were rejected, and time intervals with extreme deflection or frequency bursts were removed after visual inspection. Additionally, PCA and ICA was applied for further artifact rejection. The reader can consult (Babayan et al., 2019) for details.

#### 2.1.2 Synthetic EEG data from a population of weakly coupled oscillators

Synthetic EEG signals were created using the order parameter of a population of weakly coupled nonlinear oscillators following the Matthews-Mirollo-Strogatz model(Matthews et al., 1991). Simulations were implemented using the DifferentialEquations.jl package (Rackauckas and Nie, 2017). A system of 800 oscillators was instantiated, whose natural frequencies followed the uniform distribution with a bandwidth Δ = 0.8. Solutions for the order parameter were obtained by a fourth order Runge-Kutta method(timestep=1/250) using three distinct coupling constant *K*: 1.1(phase-locking), 0.1(incoherence), and 0.9(chaos). A24[s] epoch was isolated from each case, and white noise and pink noise were added to reproduce the normal EEG spectral background.

#### 2.1.3 iEEG data

The “atlas of the normal intracranial electroencephalogram” (AN-iEEG) dataset (Frauscher et al., 2018) was also analyzed to further study the human alpha rhythm generator dynamics. The AN-iEEG dataset is a massive resting-state iEEG dataset collected from candidates for epilepsy surgery at three different hospitals in Canada and France. The recordings were obtained while the subjects were awake with their eyes closed and were selected to exclude epileptic activity. The signals were recorded using either sEEG, subdural strips, or subdural grids at the different hospitals, and the electrode locations were projected into a common brain map. Segments 60[s] long were selected visually from each recording. The data was filtered at 0.5-80[Hz] with a FIR filter and decimated to 200[Hz]. An additional adaptive filter was used to reduce power-line noise. We only analyzed signals coming from the occipital lobe, and thus we included the following structures (number of individuals in parenthesis): superior and middle occipital gyri (21); inferior occipital gyrus and occipital pole (23); cuneus (19); calcarine cortex (12); lingual gyrus and occipital fusiform gyrus (29). See (Frauscher et al., 2018) for extra details.

### 2.2 EEG data analysis

We followed the pipeline reported by Díaz et al. (2018). All computational procedures were implemented using the Julia programming language (Bezanson et al., 2017) (https://julialang.org). From the preprocessed data, data from two individuals were discarded as time interval rejection produced too short, highly discontinuous recordings. From this group of 201 individuals, only the channels O1, Oz, and O2 from each subject were used for further manipulation. Channel length was 474.83±9.30[s] for the eyes-closed condition and 469.44±13.20[s] for the eyes-open condition.

The data was digitally filtered in the forward and backward direction with a second order Butterworth filter to isolate the alpha band. This approach effectively obtains a zero phase response and contributes to conserve the signal and its envelope waveform. Each channel was divided into 24[s] epochs (6000 points per epoch) with an overlap of 50%. This amounts to 22893 epochs for the eyes-closed condition and 22634 for the eyes-open condition. The Hilbert transform was computed for each epoch and its envelope obtained using the relation 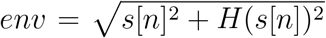. Since the envelope computation results in spurious effects at both ends, 2[s] were excised from both tails of the estimated envelope. From these 20[s] envelopes, the coefficient of variation of the envelope *CVE* = *σ_env_/μ_env_* and the mean of the envelope(ME) were estimated. CVE values were color encoded using the “buda” color scheme(Crameri, 2018) as low-CVE/purple, mid-CVE/salmon, and high-CVE/gold.

#### 2.2.1 Surrogate Data for Gaussianity

To obtain a probability model for Gaussianity, 10^6^ instances of a 6000-point vector were simulated, sampling each point from a normal standard distribution 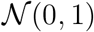. As in the experimental data, a sampling frequency of 250[Hz] was assumed. These *in silico* generated epochs were treated as previously described in subsection 2.2 for the experimental data. The same filter was applied to each epoch to obtain the alpha band. The envelope was calculated, 2[s] segments were cut from both ends, and the CVE and ME values were computed. Then, the probability density function(PDF) and the cumulative density function (CDF) were calculated for the CVE values of these alpha-filtered white Gaussian noise epochs. The 0.005 and 0.995 quantiles were estimated to obtain the lower and upper limits of a 99% confidence interval to assess the null hypothesis *H*_0_: an EEG epoch is indistinguishable from white Gaussian noise.

#### 2.2.2 Envelope characterisation space. Scatterplots, density plots and vector field

The envelope characterization space is a phase space that includes CVE and EEG amplitude measured for each epoch. EEG amplitude is represented by the mean of the envelope(ME), which is log-transformed and normalized using each subject and each subject’s channel statistics. This phase space is visualized by scatterplots and density plots. The density plots were built estimating 2D-histograms with a 500×500 matrix. Rows and columns were filtered with a 51-coefficient binomial kernel, and the matrix was plotted using an alternating colour/white palette to produce a contour-like plot. In addition, as the (CVE,log(ME)[z-scored]) data for each subject and each channel’s recording formed an independent data cloud containing both eyes-open and eyes-closed points, it is possible to use the centroid of eyes-open and eyes-closed points to analyze the overall behavior in the envelope characterization space for each experimental condition. Thus, the centroid of all channels of each subject’s data — (CVE,log(ME)[z-scored]) — was estimated for eyes-open and eyes-closed points to evaluate the change between experimental conditions. Then, a single vector was traced from the eyes-open centroid (white dot) to the eyes-closed centroid (arrowhead). This procedure results in a vector field inside the envelope characterization space, where each single vector was categorized as having low-CVE (purple), mid-CVE (salmon) or high-CVE (gold) based on the CVE coordinate of the eyes-closed centroid.

#### 2.2.3 Time-frequency analysis

Time-frequency analysis was applied to a complete recording that did not suffer any time interval removal(960[s]). The eyes-closed and eyes-open blocks were re-concatenated to recover the initial interleaved protocol. Then, the data was divided in 20[s] epochs with an overlap of 50% and treated as in Prerau et al. (2017) setting the time-bandwidth=10[s] and number of tapers=20, resulting in a frequency resolution of 0.5[Hz]. Only frequencies 20>f> 1 were kept and the spectral power was log-transformed and colour encoded (dark blue < cyan cyellow < dark red).

#### 2.2.4 Comparison of Fourier analysis and envelope analysis

One raw representative epoch for each CVE class was selected and re-analyzed using both Fourier analysis and envelope analysis. The spectrum for each epoch was calculated before filtering, while after filtering the CVE and energy were obtained and visualized in the envelope characterization space. The same steps were applied to the synthetic EEG data from a population of weakly coupled nonlinear oscillators.

### 2.3 iEEG data analysis

The iEEG data was treated as the EEG data, with some exceptions. The CVE surrogate distribution was re-calculated setting the sampling frequency to 200[Hz] to match the AN-iEEG sampling frequency. Filtering, segmentation, and envelope-parameter computation remained untouched. In total, 416 epochs were analyzed. In the envelope characterization space, energy values were log-transformed but not normalized, as the AN-iEEG dataset contains eyes-closed data only. Three representative epochs recorded from the calcarine cortex were used to compare Fourier analysis with envelope analysis.

## 3 Results

### 3.1 EEG envelope analysis

#### 3.1.1 CVE for alpha-filtered Gaussian noise

The main motivation behind the envelope analysis rests upon the statistical characteristic of Gaussian signals and their envelopes. Therefore, we first review these mathematical foundations. For any zero-mean continuous-time Gaussian signal (and any of its filtered sub-bands) (Fig. 1.a), the respective envelope signal follows a Rayleigh distribution (Fig. 1.b). Since the coefficient of variation for the entire family of Rayleigh distributions is invariant and equal to 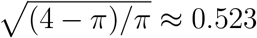, the coefficient of variation of the envelope (CVE) of a infinite zero-mean Gaussian signal always display this value that acts as a fingerprint for Gaussianity (Fig. 1.c). Thus, CVE is a dimensionless, scale-independent parameter to assess Gaussianity. Nonetheless, 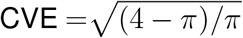 only holds for infinite signals, and the CVE distribution for discrete-time Gaussian signals must be estimated by computer simulations. This distribution varies both with epoch length and bandwidth (as well as sampling frequency).

**Figure 1:**
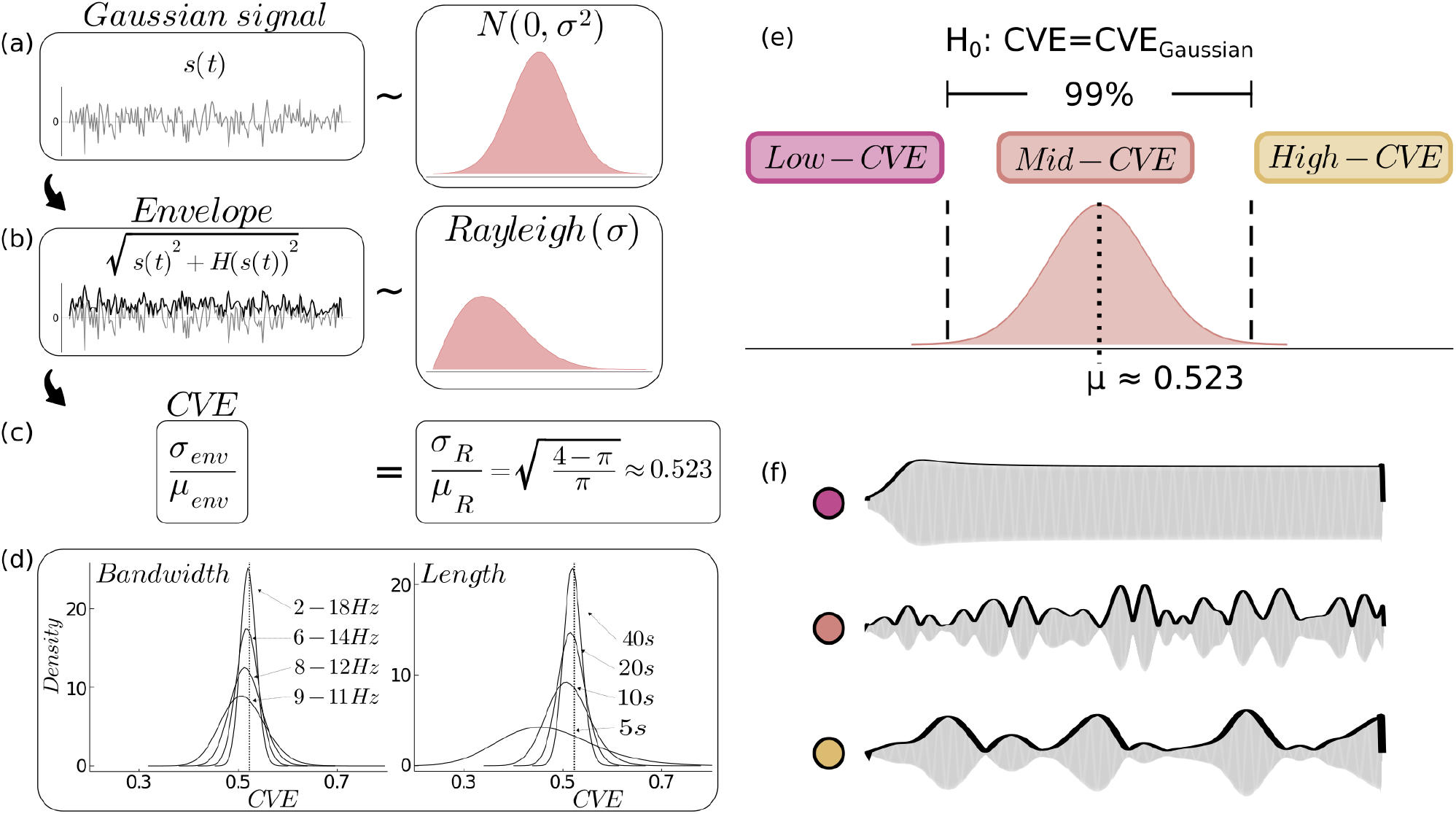
Coefficient of variation of the envelope is a fingerprint of Gaussianity. **(a)** The continuous-time Gaussian noise signal s(t) follows a Gaussian distribution N ~ (0, σ^2^). **(b)** The envelope env(t) is the norm of the analytic signal z(t) = s(t) + iH(s(t)), where H(s(t)) is the Hilbert transform of s(t). Thus, 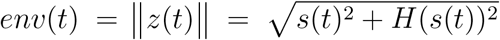. The envelope of s(t) follows the Rayleigh distribution. **(c)** The coefficient of variation of the envelope (CVE) of s(t) is the coefficient of variation of the Rayleigh distribution. This value is invariant and equal to 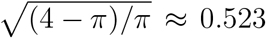 **(d)** Since the identity 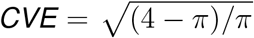 only holds for infinite, continuous-time signals, the CVE distribution for discrete-time Gaussian noise epochs of arbitrary length is obtained by computer simulation. CVE distributions for 24[s] epochs filtered at different bandwidths (left panel) show a mode 0.523 (dotted line) and a variance → 0 as the bandwidth →+∞. CVE distributions for arbitrary length epochs filtered between 7-14 (right panel) show the same trend. The mode → 0.523 and the variance → 0 as the epoch length → +∞. All epochs were sampled at 250[Hz]. Notice these distributions also vary with the sampling frequency(not shown). **(e)** The CVE distribution for discrete-time Gaussian noise allows constructing confidence intervals (both dashed lines) to assess the null hypothesis of Gaussianity. Hence, with respect to CVE, three different classes of behavior can be established: low-CVE (purple), mid-CVE (salmon), and high-CVE (gold). **(f)** These CVE classes correspond to distinctive amplitude modulation patterns that are linked to different generative processes. Low-CVE signals are synchronous, rhythmic signals coming from phase-locking processes, while mid-CVE signals correspond to Gaussian noise, and high-CVE signals are distinctively non-sinusoidal and pulsating.

We first estimated the CVE probability density functions for Gaussian signals of different lengths (5,10,20,40 [s]) and bandwidths(2,4,8,16 [Hz]) sampled at 250[Hz](Fig. 1.d). All distribution are bell-shaped with a mode close to the CV of the Rayleigh distribution and their variance decreases as the epoch length and the bandwidth increase. An equivalent trend is obtained when varying the sampling frequency (not shown). Thus, the surrogate CVE distribution allows to establish a confidence interval for the null hypothesis *H*_0_: epoch indistinguishable from Gaussian noise. As a consequence, three different CVE classes are also established: low-CVE, mid-CVE, and high-CVE (Fig. 1.e). Additionally, each CVE class displays distinctive signal morphologies produced by different amplitude modulation patterns, and thus reflect distinct underlying generative processes (Fig. 1.f).

#### 3.1.2 Alpha-band CVE confidence interval and CVE for resting state EEG

To characterize the human alpha rhythm using the CVE, we first obtained confidence intervals of Gaussianity to test the experimental resting-state EEG epochs from the LEMON dataset. For our simulations, we used 24[s] epochs with a passband in ≈ 8-13 following the definition of Kane et al. (2017). As expected, the CVE probability density function for alpha-filtered Gaussian noise epochs obtained by simulations is bell-shaped and unimodal with mean= 0.520, which is near the fingerprint of Gaussianity 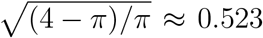 (Fig. 2, gray trace, both panels). The 0.005 and 0.995 quantiles in this distribution are 0.460 and 0.586, respectively. These values define the 99 % confidence interval for our hypothesis of Gaussianity and establish three CVE classes: low (CVE<0.460), mid (0.460< CVE <0.586) and high (0.586<CVE). Thus, any epoch with a CVE value inside the interval [0.460,0.586] is considered to be indistinguishable from Gaussian noise. The CVE distributions for eyes-open and eyes-closed experimental data is shown in Fig. 2. While both distributions exhibit positive skewness and have their medians and modes inside the Gaussian region, they differ in their relative contribution of each CVE class. Eyes-open data (right panel, black trace) display values almost exclusively in the mid-CVE and high-CVE intervals. On the other hand, eyes-closed data (left panel, black trace) display values in all three CVE intervals, with ≈ 10% of its values in the low-CVE region.

**Figure 2:**
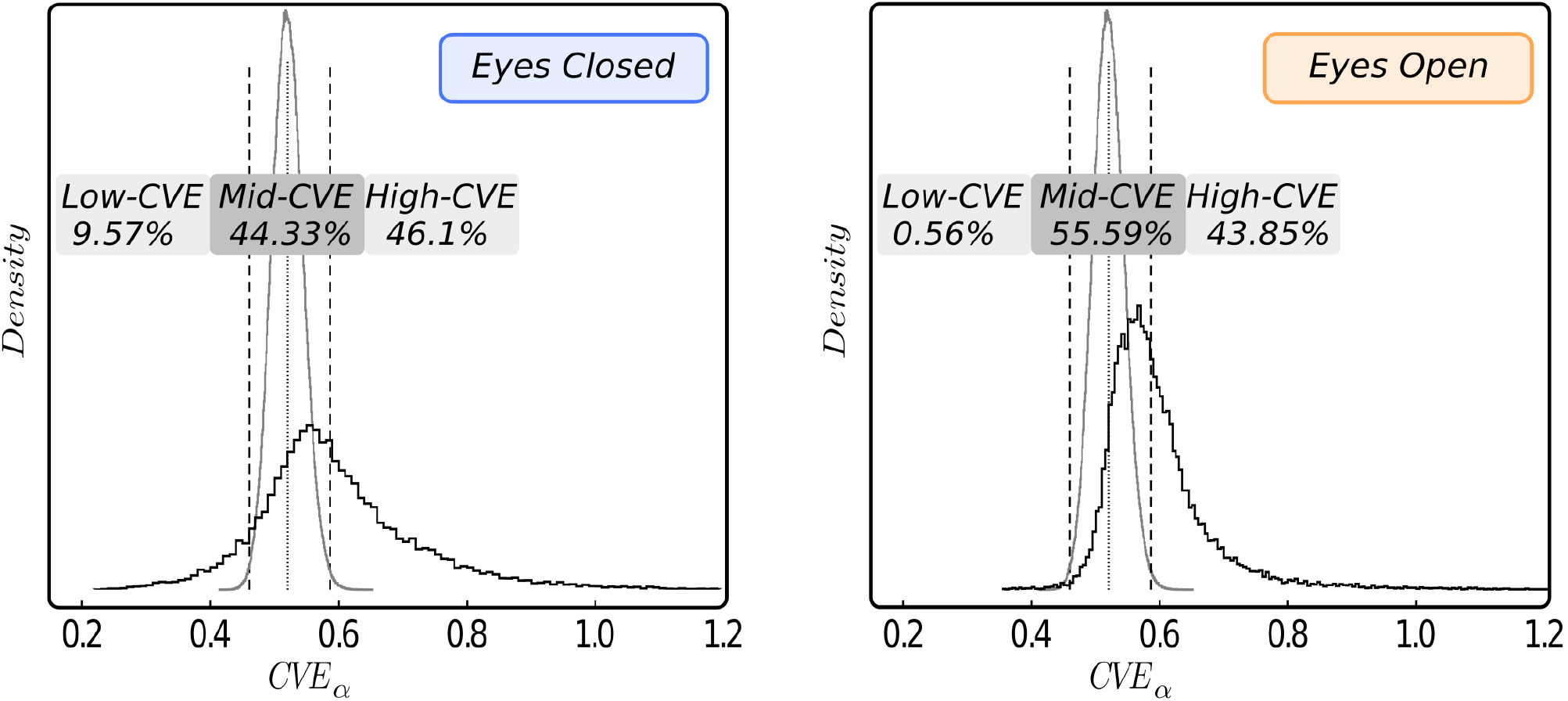
CVE distribution for surrogate and LEMON dataset experimental data. The CVE distribution of alpha-filtered Gaussian noise simulated epochs (grey trace, both panels) is unimodal and bell-shaped with a mean equal to 0.520 (dotted line). Dashed lines at 0.460 and 0.586 depict the 99 % confidence interval for Gaussianity. Both empirical distributions (black traces) for eyes-closed and eyes-open conditions are unimodal and show a mode and median inside the confidence interval for Gaussianity. The relative frequency of epochs corresponding to each one of the three CVE classes is shown as percentages above each distribution. Left panel: The eyes-closed data distribution (22893 epochs) is nearly symmetrical with long tails toward low-CVE and high-CVE values. Almost 10 % of experimental epochs have a low-CVE while 90 % are in the Mid-High CVE ranges. Right panel: The eyes-open data distribution (22634 epochs) presents most of its values in the mid-CVE region and a long tail in the high-CVE interval. Low-CVE epochs are almost absent.

#### 3.1.3 Envelope characterization space and vector field

The *envelope characterization space* is a two-dimensional phase space based on two envelope signal pa-rameters: (i) coefficient of variation of the envelope (CVE) and (ii) mean of envelope (ME). Both values have to be combined in a single framework as the CVE is a scale-independent statistic. Scatterplots in Fig. 3 show the envelope characterization space for eyes-closed and eyes-open conditions (all channels together). In both conditions, epochs with the lowest amplitude display CVE values inside the Gaussian region. While eyes-closed data (blue) exhibit high amplitude epochs in all three CVE regions, low-CVE epochs group around one standard deviation above the mean energy. On the other hand, mid-CVE and high-CVE epochs are more broadly distributed over the high energy region. Eyes-open data (orange) mainly contain low amplitude epochs (as expected from the experimental condition), displaying almost exclusively mid-CVE and high-CVE values.

**Figure 3:**
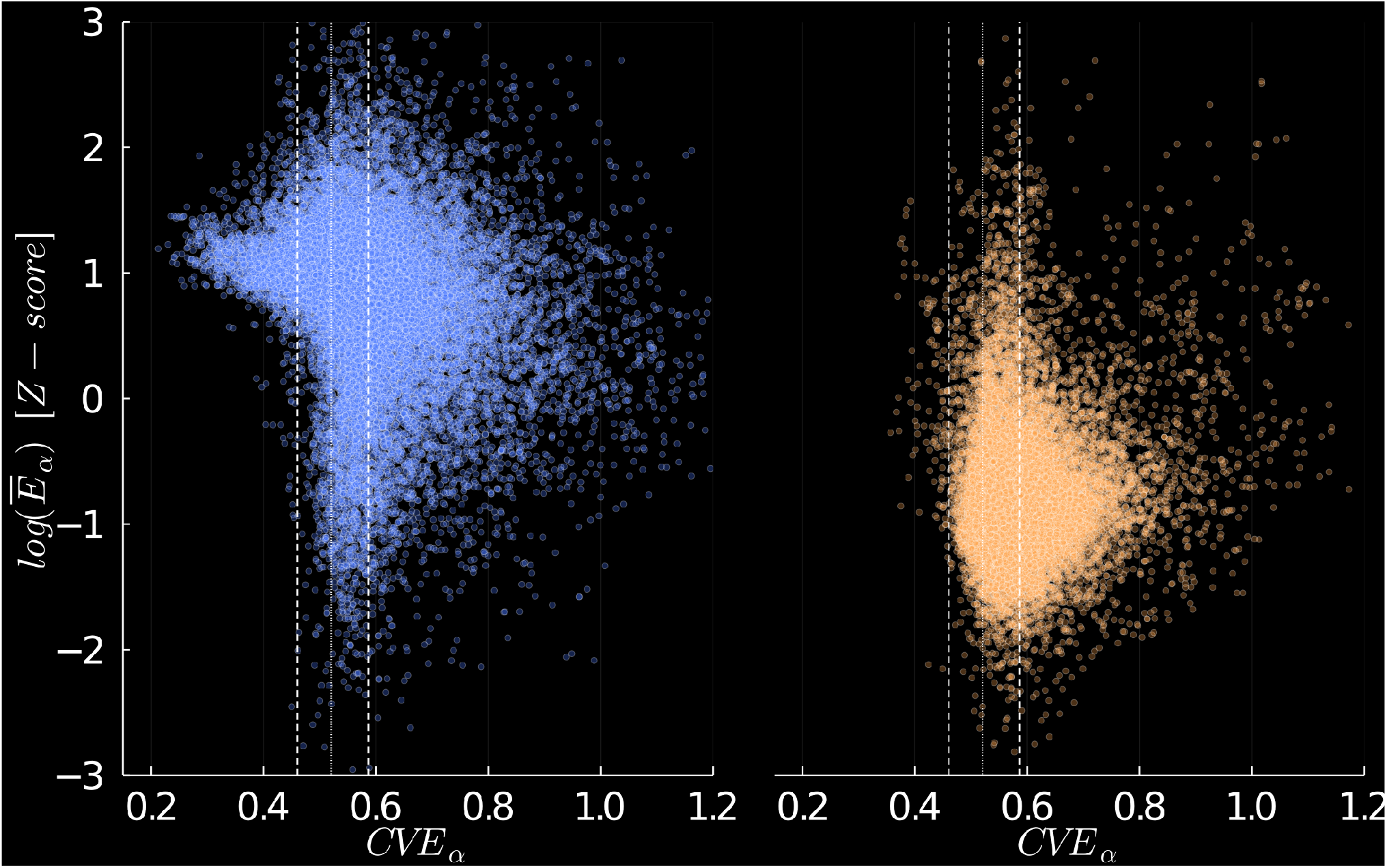
Scatterplot of EEG amplitude versus coefficient of variation of the envelope (CVE) for eyes-closed and eyes-open conditions. The total data from 201 subjects and 3 EEG channels (O1,Oz,O2) was used for this plot. Each channel data was split in eyes-closed (blue) and eyes-open data (orange), filtered in the alpha band and epoched in 24[s] segments with an overlap of 12[s]. For each corresponding epoch, the mean of the envelope (ME) and CVE were calculated. Then, the ME was log-transformed and Z-scored using the mean and standard deviation from the whole recording (eyes-closed and eyes-open), resulting in normalized envelope amplitude values for each subject and each channel data. Left panel: Eyes-closed data (22893 epochs) show a CVE excursion from 0.3 to 1.2. The epochs corresponding to low-CVE values also have an energy one standard deviation above the mean. The big majority of epochs have CVE values corresponding to Gaussian or pulsating signals. Some epochs display an energy level below the mean energy, which corresponds to subjects whose alpha rhythm is either weak or non-existent (as widely reported in the literature). Additionally, high energy epochs are present in all three CVE classes. Right panel: Eyes-open data (22634 epochs) mostly show amplitude below the mean energy, as expected from the experimental condition, and with a clear tendency towards mid-CVE and high-CVE values. Low-CVE values are almost absent. Dotted line at 0.520, the mean of the surrogate alpha-filtered Gaussian noise CVE distribution. Dashed lines at 0.460 and 0.586 depict the 99 % confidence interval for Gaussianity. Transparency was set to 50% to highlight cluster density.

Contour plots permit further analysis of the envelope characterization space (Fig. 4). Eyes-closed data (blue) strongly cluster in the Gaussian region around 1 standard deviation above the mean. While eyes-open data (orange) also clusters in the Gaussian region, most epochs group around 1 standard deviation below the mean.

**Figure 4:**
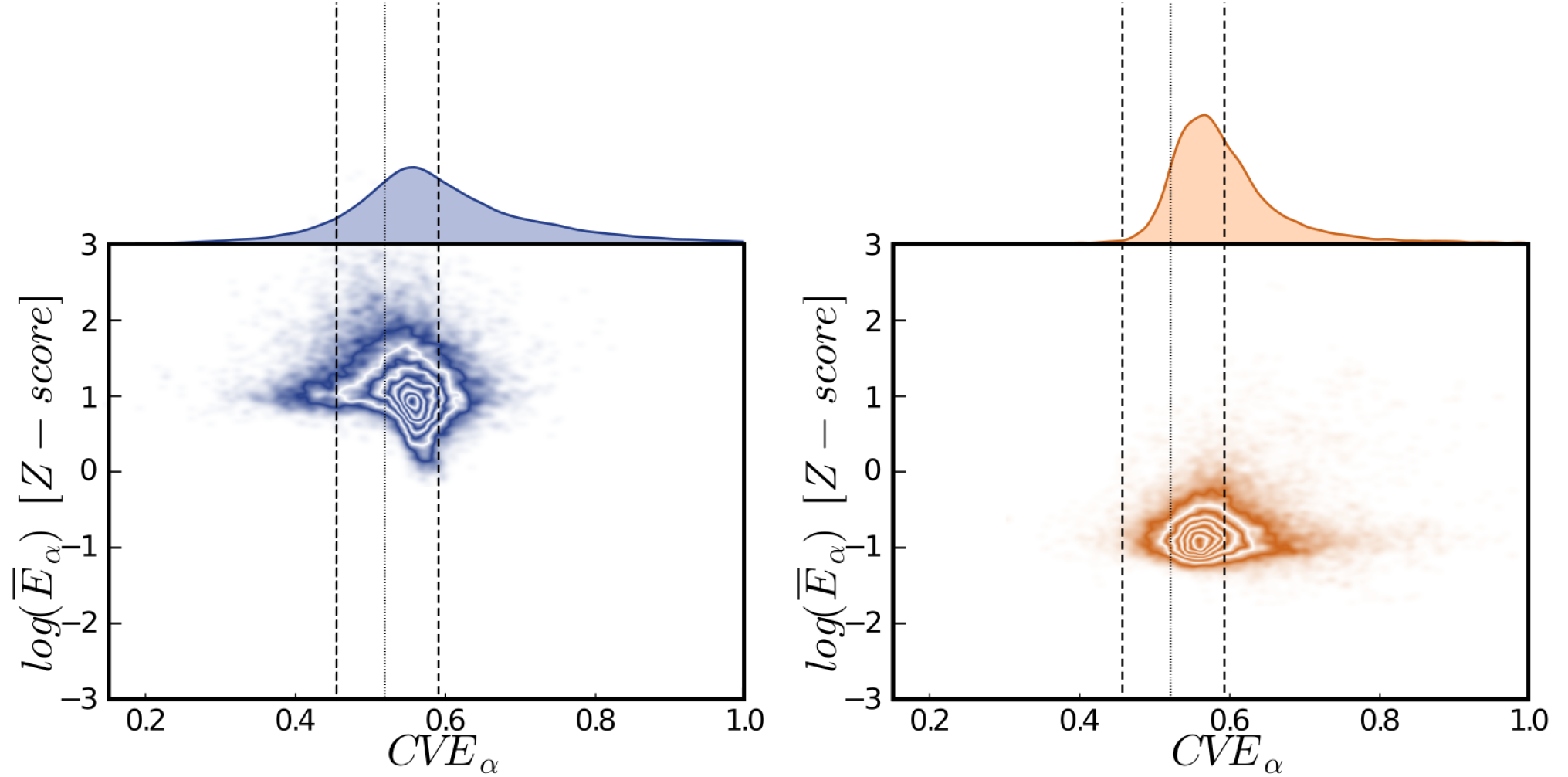
Contour plots reflecting the relationship between EEG amplitude and CVE. Each panel shows the empirical distribution of CVE (upper trace, same as in Figure 1) and a contour plot relating mean average energy (y-axis) with CVE values for each epoch. Left panel: Eyes-closed data presents one cluster in the interval of Gaussianity with an amplitude approximately 1 standard deviation above the mean. The data display CVE values in all three CVE regions. Both eyes-closed (blue) and eyes-open (orange) data present just one cluster centered at similar mid-CVE values but differ in amplitude. Right panel: Eyes-open data also show just one cluster again centered on the region of Gaussianity. This cluster, however, resides in the low energy region, approximately 1 standard deviations below the mean energy level. Dotted line at 0.520, the mean of the surrogate alpha-filtered Gaussian noise CVE distribution. Dashed lines at 0.460 and 0.586 depict the 99 °% confidence interval for Gaussianity.

This difference of about 2 standard deviations in the overall amplitude change between both conditions is also found when data from each subject and each channel are analyzed independently. Vectors from the eyes-open centroid to the eyes-closed centroid were traced and plotted in the envelope characterization space (Fig. 5). Each of these vectors was classified as low-CVE, mid-CVE or high-CVE based on the CVE coordinate of the eyes-closed centroid. While there is some variability in the magnitude and phase of these vectors(especially high-CVE vectors), the overall trend is an amplitude change of about 2 standard deviations in all three classes of vectors. The presence of low-CVE vectors shows that some subjects generate preferentially rhythmic alpha rhythms, although many subjects generate data that is either Gaussian or pulsating overall.

**Figure 5:**
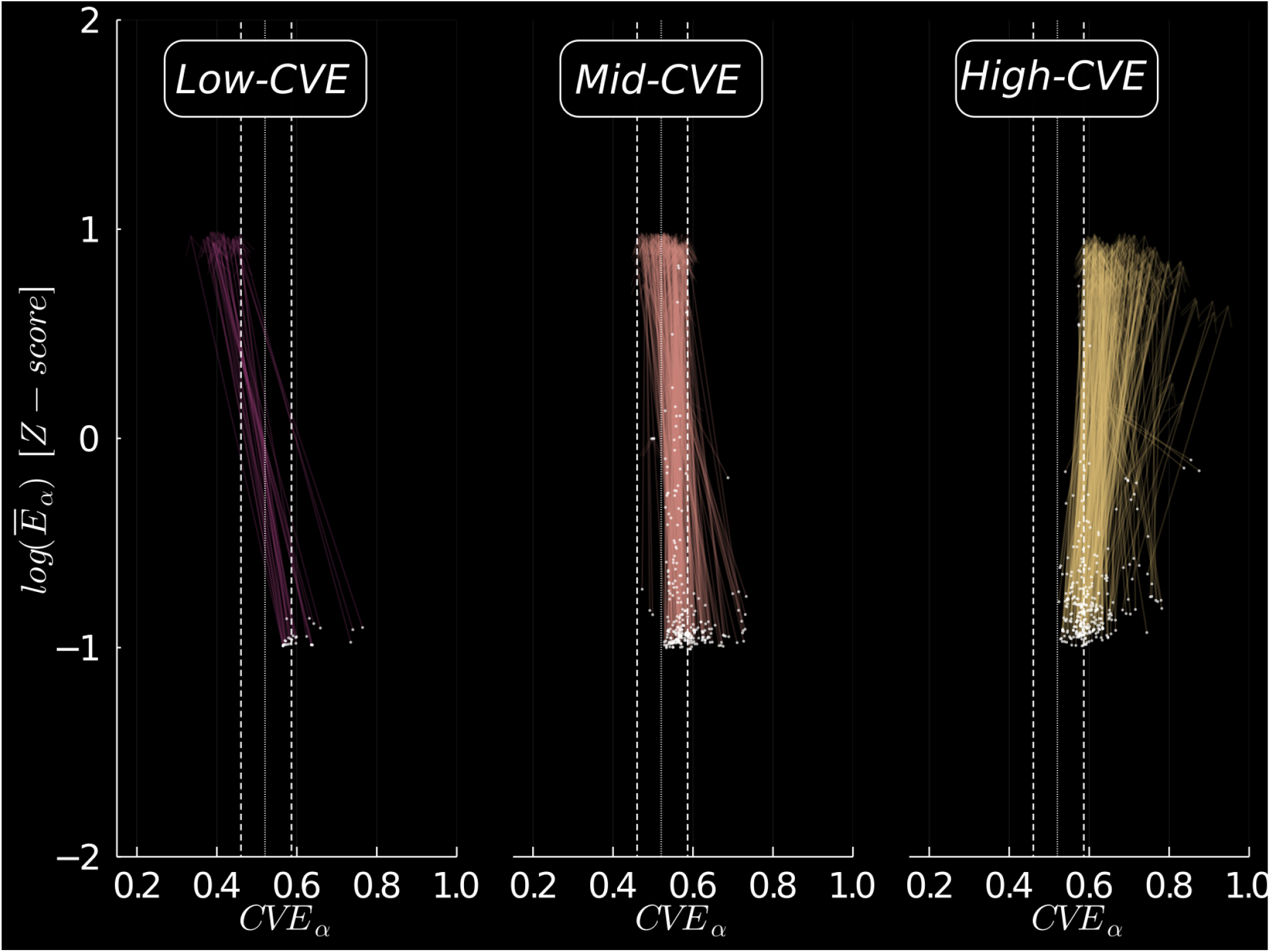
Vector field in the envelope characterization space for eyes-open to eyes-closed transitions. The centroid of CVE and energy values from each subject (N=201), each channel (O1,Oz,O2) data were calculated. A single vector was traced starting from the eyes-open centroid (white dot) to the eyes-closed centroid (arrowhead). Vectors were classified as either low-CVE, mid-CVE or high-CVE according to the CVE coordinate of their end position (eyes-closed centroid), reflecting the overall behavior of subject and EEG channel specific data. While almost all vectors start at mid-CVE and high-CVE positions and in the low energy range, their endpoints finish in all three CVE intervals and with an energy gain of 2 standard deviations. Most transitions end up in CVE regions that do not reflect rhythms (Mid-CVE and High-CVE) and the relative frequency of vectors endpoint roughly reflects the entire population CVE distribution. Overall, the great majority of transitions starts as a Gaussian like profile and dot not end as a clear a like oscillation. Dotted line at 0.520, the mean of the surrogate alpha-filtered Gaussian noise CVE distribution. Dashed lines at 0.460 and 0.586 depict the 99 °% confidence interval for Gaussianity. The transparency channel was set to 50% to highlight cluster density.

#### 3.1.4 Spectrogram and CVE during open/closed conditions

The spectrogram of a complete recording (960[s]) shows that alpha power increases during the eyes-closed condition regardless of the CVE values involved (Fig. 6, left panel), as CVE is straddles on all three CVE intervals. During the eyes-open condition, CVE tends to settle inside the Gaussian region. Spurious high-CVE epochs are often found in the transition between conditions as those epochs contain both eyes-open (low energy) and eyes-closed (high energy) data, thus appearing pulsating. Three representative eyes-closed epochs associated with high alpha power display the morphology/amplitude modulation characteristic of their respective CVE values (Fig. 6, right panel).

**Figure 6:**
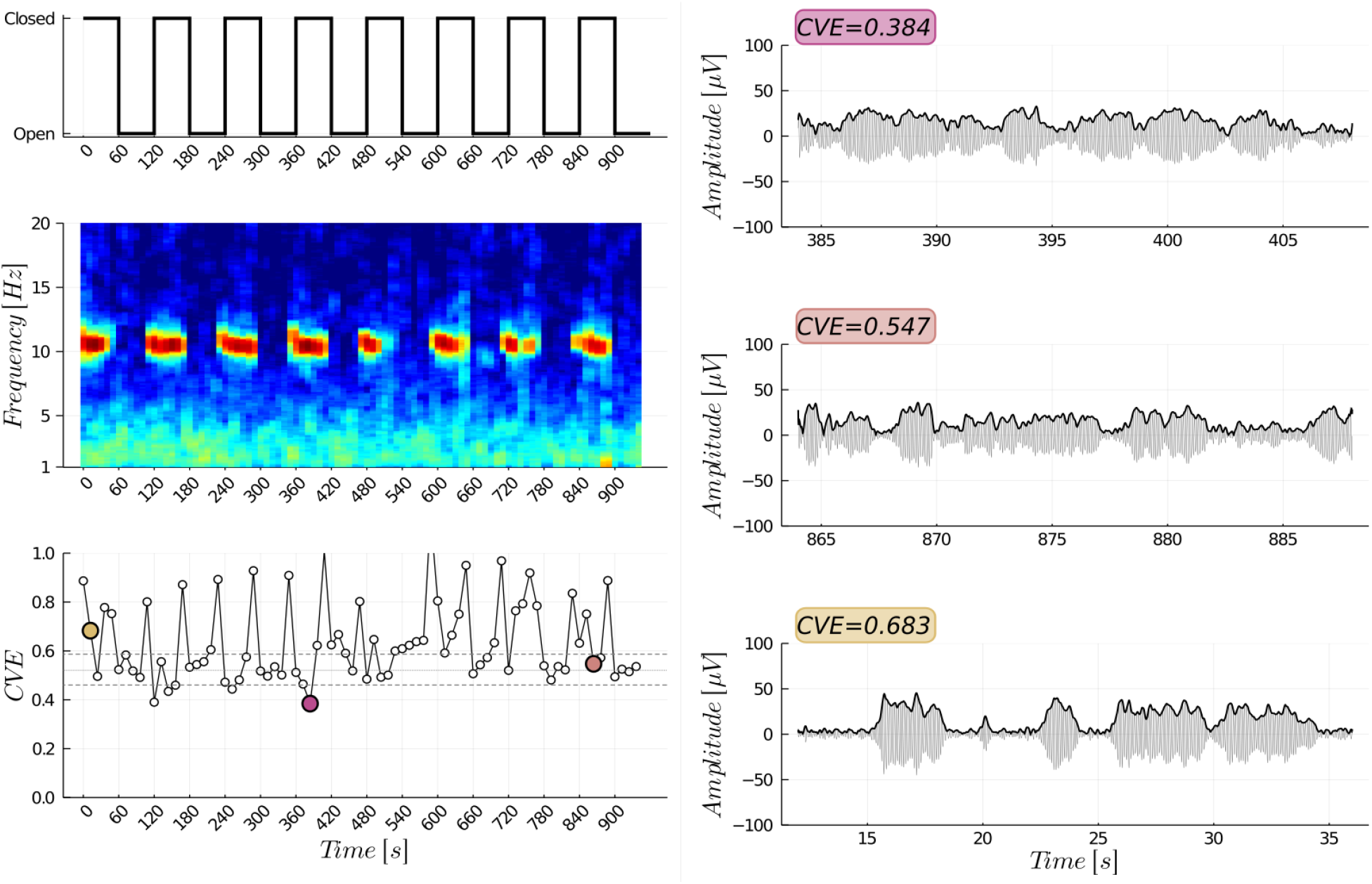
Relation between experimental condition, EEG time-frequency analysis and CVE temporal changes. Data correspond to a complete 960[s] recording (channel O1) from a representative individual. Left panel: from top to bottom, experimental condition, multitaper spectrogram (data not filtered) and CVE temporal profile (alpha-filtered data). While both conditions show a broadband 1/f-like spectrum, only the eyes-closed condition contain a narrowband component ≈ 10[Hz]. Furthermore, the CVE shows low, mid and high CVE values associated with the narrowband spectrum during the eyes-closed condition. On the other hand, only mid-CVE and high-CVE values are present during the eyes-open condition. The right panel depicts three eyes-closed 24[s] epochs (color-coded according to the data points highlighted in the CVE profile) with their respective envelopes corresponding to rhythmic (upper trace), Gaussian (middle trace) and pulsating (lower trace) activity. Dotted line at 0.520, the mean of the surrogate alpha-filtered Gaussian noise CVE distribution. Dashed lines at 0.460 and 0.586 depict the 99 % confidence interval for Gaussianity.

#### 3.1.5 Fourier/envelope analysis comparison

As some authors have argued in favour of time domain methods over frequency domain methods to study brain rhythms (Jones, 2016), we re-analyzed three representative epochs (same data in Fig. 6) using Fourier analysis and envelope analysis (Fig. 7). A closer inspection of their pre-filtering spectra reveals that all three epochs contain the canonical spectral peak at « 10[Hz]. Furthermore, their spectral profiles are almost indistinguishable from each other (Fig. 7.b). The CVE, however, can easily discriminate among these epochs, as shown in the envelope characterization space (Fig. 7.c). A similar trend was obtained through analysis of synthetic signals obtained from the order parameter of a synchronization MMS model of coupled oscillators ((Matthews et al., 1991)). While the three alpha-filtered epochs display different morphologies (Fig. 7.d), their pre-filtering spectra cannot be separated from each other (Fig. 7.e). Noticeably, the spectral peak at « 10[Hz] is much narrower than for experimental data. Again, these three synthetic epochs can be distinguished by their CVE in the envelope characterization space (Fig. 7.f).

**Figure 7:**
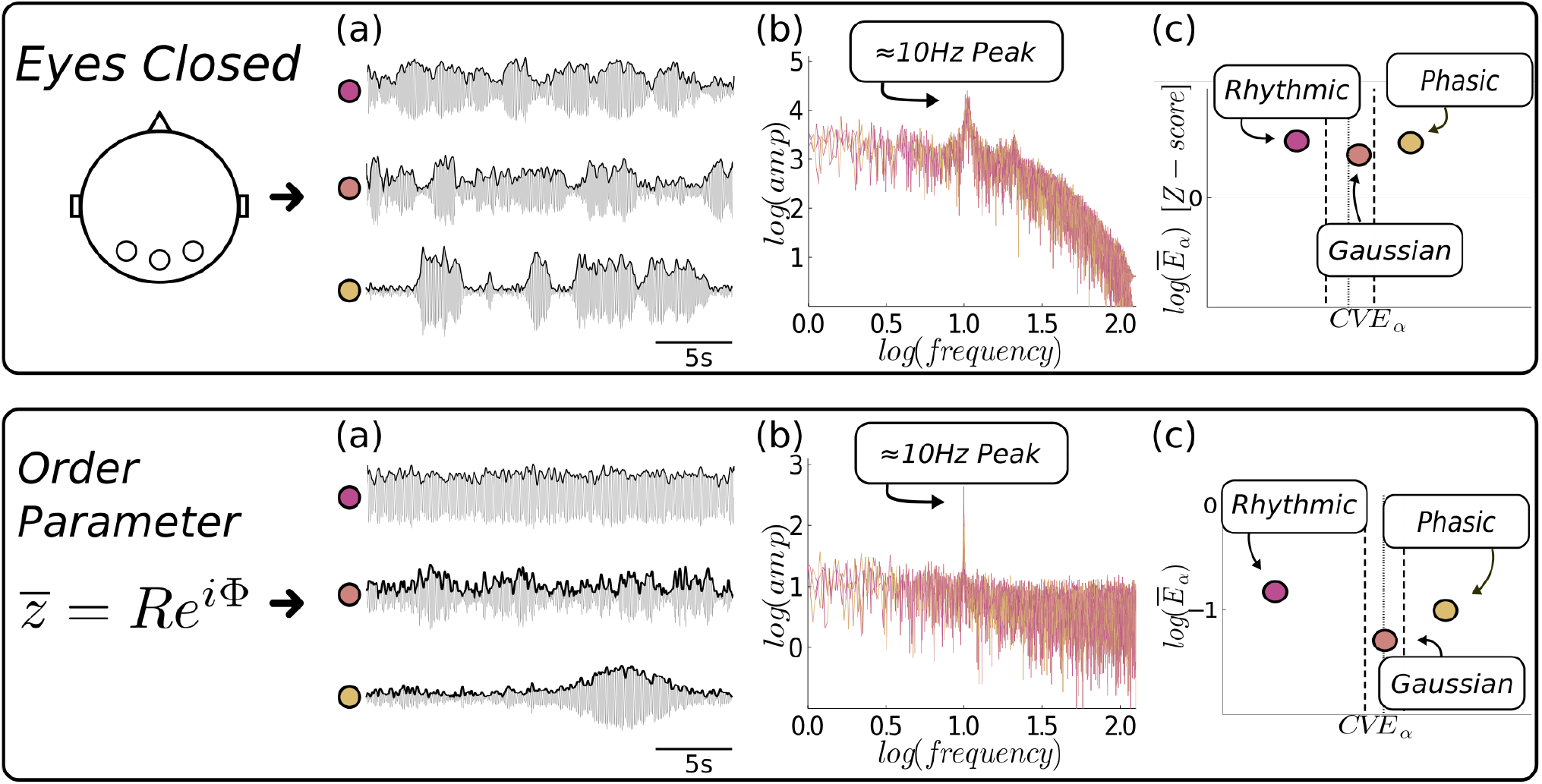
Envelope analysis reveals both Gaussian and synchronous dynamics in event-related activity. Three eyes-closed EEG epochs were re-analysed using Fourier analysis(raw data) and envelope analysis(alpha-filtered data). **(a)** Three experimental alpha-filtered epochs and their respective envelopes are shown(same data in Fig. 6), each epoch belongs to one CVE class (rhythmic, Gaussian, phasic). **(b)** The three epochs contain an energy peak ≈ 10[Hz] and their spectra are almost indistinguishable from each other, **(c)** On the other hand, the envelope analysis for the alpha-filtered data does discriminate between the three epochs by CVE. **(d)**, **(e)**, **(f)** Data obtained from the synchronization model (MMS, (Matthews et al., 1991)). Epochs with different CVE, but similar spectra and showing a marked peak at 10 Hz but the epochs can be separated by their CVE in the envelope characterization space. Dotted line at 0.520, the mean of the surrogate alpha-filtered Gaussian noise CVE distribution. Dashed lines at 0.460 and 0.586 depict the 99 % confidence interval for Gaussianity.

### 3.2 Envelope analysis of intracranial EEG data

Epochs obtained from occipital intra-cortical EEG data (AN-iEEG dataset) are highly clustered in the Gaussian region, while only a small percentage of epochs (5%) stretch into the low-CVE region (Fig. 8). We also analyzed AN-iEEG data specifically recorded from the calcarine cortex (Fig. 8, right panel), a neuroanatomical region that Lopes Da Silva and Storm Van Leeuwen (1977) suggests as a possible generator for the alpha rhythm. Similarly, epochs recorded from the calcarine cortex exclusively contain CVE values indicating Gaussian EEG segments, the percentage of synchronous epochs (i.e. having a low-CVE) is negligible.

**Figure 8:**
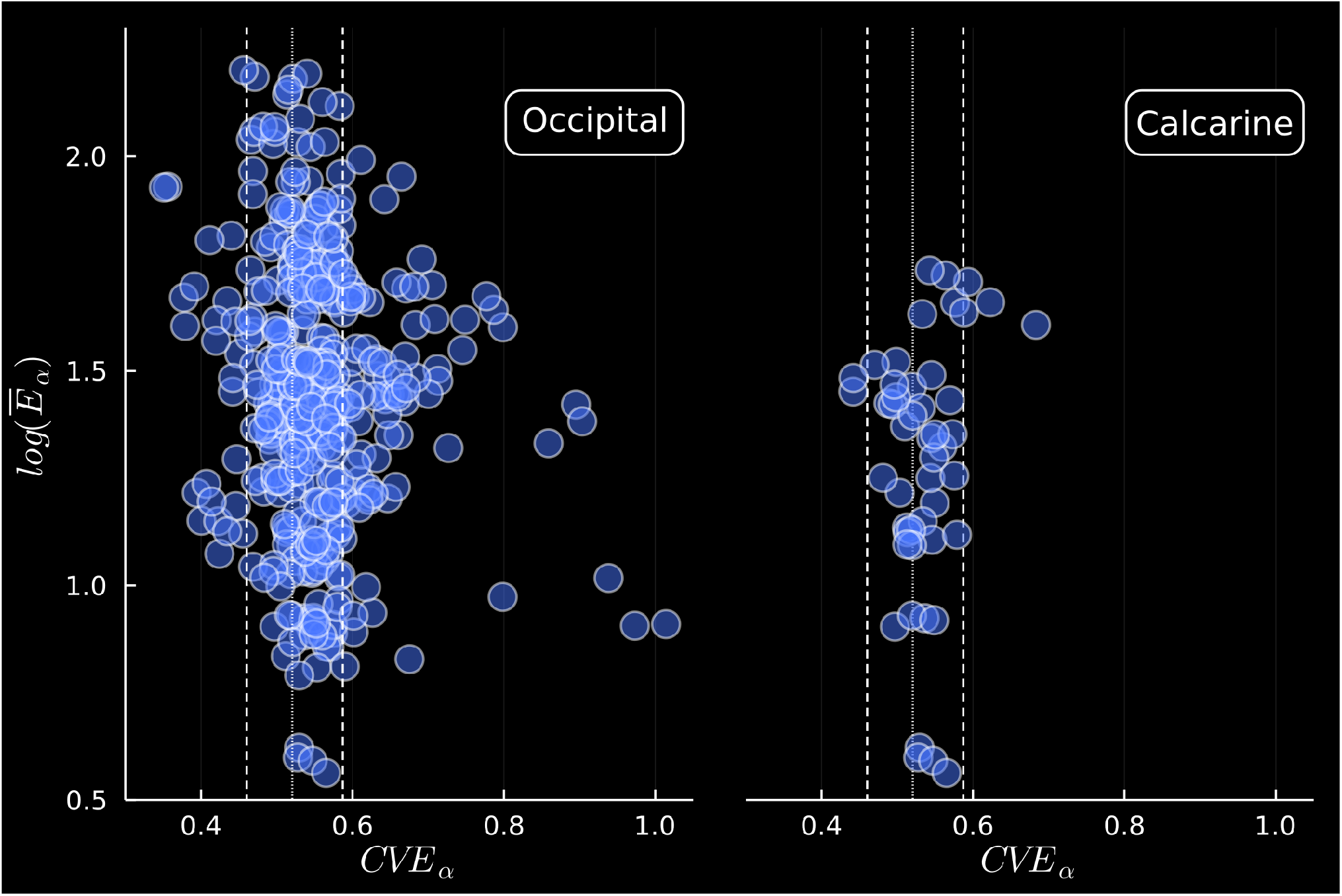
Scatterplot of epoch amplitude versus coefficient of variation of the envelope (CVE) for AN-iEEG dataset recordings obtained under eyes-closed condition. The recordings were filtered in the alpha band and epoched in 24[s] segments with an overlap of 12[s]. The mean of the envelope (ME) and CVE were calculated, and the ME was log-transformed. Left panel: Occipital data (412 epochs) is highly clustered in the Gaussian region, while 24 epochs lie in the low-CVE (rythmic) region. Right panel: Data subset obtained only from the Calcarine cortex (48 epochs) mostly contains Gaussian epochs and synchronous epochs are practically absent. Dotted line at 0.520, the mean of the alpha-filtered Gaussian noise CVE distribution obtained by computer simulations. Dashed lines at 0.455 and 0.592 depict the 99 % interval for H_0_: epoch indistinguishable from Gaussian noise.

We applied Fourier analysis and envelope analysis to calcarine cortex data and found the same result as when we analyzed experimental EEG and synthetic data (Fig. 9). The spectral profile for calcarine epochs is almost the same for all, and the spectral peak at « 10[Hz] is much broader than the scalp-level EEG and synthetic counterparts (Fig. 9.b). Nonetheless, these three iEEG epochs display different amplitude modulation regimes; a feature that is revealed in their CVE (Fig. 9.c).

**Figure 9:**
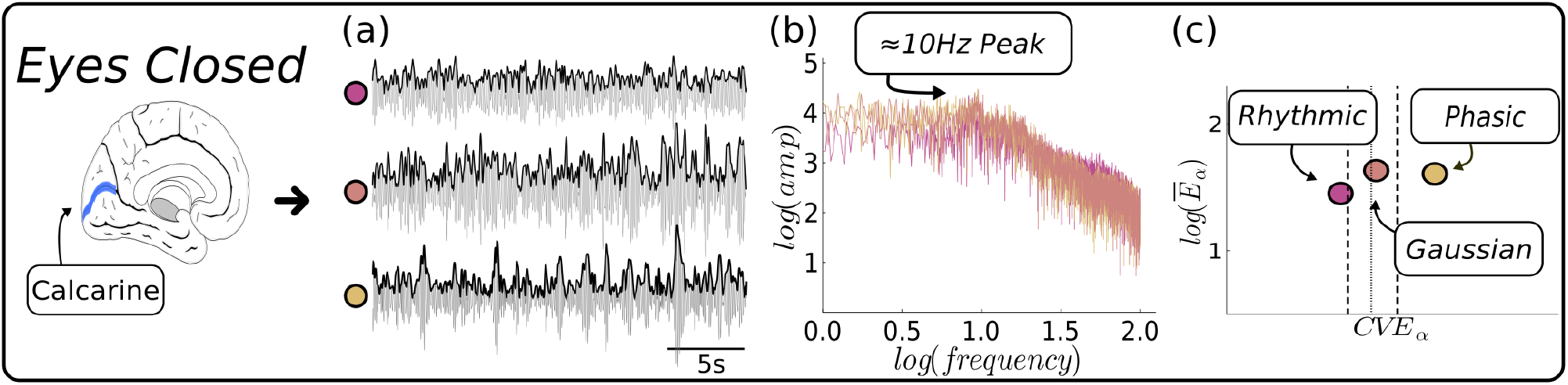
Envelope analysis reveals both Gaussian and synchronous dynamics in intracranial event-related activity in the alpha band. Three eyes-closed AN-iEEG dataset epochs recorded from the calcarine cortex were re-analysed using Fourier analysis (raw data) and envelope analysis (alpha-filtered data). **(a)** Three experimental alpha-filtered epochs and their respective envelopes are shown, each epoch corresponding to one CVE class (rhythmic, Gaussian, phasic). **(b)** The three epochs contain energy at ≈ 10[Hz] and their spectra are very similar. The spectral peak is much broader than the observed in the EEG or synthetic data (Fig. 7). **(c)** Although their spectra are similar, the envelope analysis for the alpha-filtered data discriminates between the three epochs by CVE in the envelope characterization space. Dotted line at 0.520, the mean of the surrogate alpha-filtered Gaussian noise CVE distribution. Dashed lines at 0.460 and 0.586 depict the 99 % confidence interval for Gaussianity.

## 4 Discussion

In this study we used envelope analysis to study the human alpha rhythm. The core of our method is the observation that, for Gaussian noise (or any its filtered sub-bands), the coefficient of variation of the envelope adopts the value 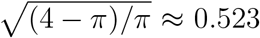 independently of the mean power of the EEG epoch under study. We demonstrated in Figures 1 and 2 the basic statistical ideas needed to work with the CVE of a given signal and how to build confidence intervals by computer simulations for the hypothesis that a given EEG epoch is indistinguishable from Gaussian noise. Furthermore, we introduced the concept of the envelope characterization space as a method to classify EEG epochs according to their mean energy and their CVE. Interestingly, in this phase space, the data clouds derived from eyes-open and eyes-closed condition are very different. Our results underscored the classic result that EEG epochs for eyes-closed are more energetic. We found (Figure 3 and 4) that most epochs for the eye-closed condition are similar to a Gaussian signal (i.e signal that looks like noise) or have a phasic temporal profile. Also in the envelope characterization space, we quantified the mean transitions between eyes-open and eyes-closed conditions (Figure 5) and found that most transitions begin as a random noise and end up as either random noise or pulsating activity. An important observation is that signals with similar frequency contents could have different CVE (Figure 6 and 7). The envelope analysis is not restricted to scalp EEG signals. In effect, iEEG signals can be classified using their CVE (Figure 8) and even signals recorded from the Calcarine cortex (visual cortex) show the same dynamics where epochs with similar spectra can be distinguished by their CVE.

Taken together, our results open another avenue for analyzing brain signals in addition to the classic use of power spectra. By itself this result is important as it provides a new heuristic tool to classify neural signals, expanding upon the use of the Fourier spectra.

However, our results not only provide a technical advantage because they open a new viewpoint to understand neuronal dynamics, but they also expand on Lord Adrian’s concept of neural synchronization. In his view, the generation of a well-defined rhythm is due to the common beat of many neurons that, via phase synchronization, generate the macroscopic *α* wave. But the CVE and the early intuition from Motokawa in 1943 show that another interpretation is possible. Motokawa approximated the Rayleigh distribution from human alpha rhythm signals in the context of his statistical mechanics theory of EEG. Thus, an epoch that has a maximal frequency content around 10[Hz] could appear solely due to the random combination of individual oscillators. This statement is true for EEG and artificial epochs (Figure 7) as well as signals recorded in the supposed cradle of *α* rhythm (Figure 9). Interestingly, most EEG epochs analyzed in this work carried the hallmark of Gaussianity, thus supporting Motokawa’s interpretation of asynchrony. However, an important number of epochs were found to be highly rhythmic, as shown by their low-CVE values, in accordance with Lord Adrian’s interpretation of neural synchronization. It seems that envelope analysis invites us to expand our viewpoints beyond the ideas of event-related synchronization (ERS) and desynchronization (ERD) and consider that the internal traffic of neural signals may be more akin to the coupling among randomly interacting oscillators.

## 5 Concluding remarks

Here, we used envelope analysis to characterize the human alpha rhythm. Envelope analysis offers an elegant time-domain method to assess Gaussianity based on the study of the envelope signal (Hidalgo et al., 2022). Previous studies regarding EEG Gaussianity can now be reassessed by envelope analysis (e.g. Elul, 1969). Furthermore, envelope analysis puts neural synchronization and Gaussianity in the framework of coupled oscillators systems (K,MMS), and it allows the continuous monitoring of neural processes purely by CVE. Our results confirm the interpretations of both Adrian (synchronization by coupling) and Motokawa (random addition of elements leading to Gaussian noise), opening a new door to explore dynamics in EEG and other neural signals (e.g. LFP, iEEG, MEG) (Rao and Edwards, 2008).

## Declaration of competing interest

None of the authors have potential conflicts of interest to be disclosed.

## Acknowledgments

This work was supported by Fondecyt (ANID Chile) 1210069 and the Japan Society for the Promotion of Science (Grant-in-Aid for Scientific Research (C): 19K12199). We are grateful for Diane Greenstein’s editorial assistance.

## Notes

### Competing Interest Statement

The authors have declared no competing interest.

